# Reduction of Weight Gain by Modulating Mitochondrial Energy Coupling Efficiency Using a Bromo-Coenzyme Q_10_ analog (6-Br-Q_0_C_10_)

**DOI:** 10.64898/2026.05.23.723575

**Authors:** Claire Yu, Wenfei Huang, Brandon Yu, Sulie L. Chang, Chang-An Yu

## Abstract

Synthetic 6-Br-Q0C10 has been shown to have a partial electron transfer activity of native coenzyme Q in the isolated mitochondria. It reduces energy coupling efficiency by 30 %, suggesting that it may be useful in the management of obesity. The effect of 6-Br-Q0C10 on cell growth has been confirmed by several cell lines. Whether or not it behaves in the same way in the animal, however, has not yet been tested. Recently, we investigated the effect of 6-Br-Q0C10 on growth of rats. When 6-Br-Q0C10 was dissolved in different media, such as carboxymethyl cellulose, ethanol, and mixture of oil and butter and then feed to rats It shows no toxicity and little negative effects on growth as measured body weight gains over a period of time. When higher concentration (0.5 mg) of 6-Br-Q0C10 was given to each rat in 0.3 mL of oil/butter (70%/30%) mixture via intragastric injection daily for a period, a significant reduction in body weight gains was observed. These results validate the earlier observation that 6-Br-Q0C10 reduces the growth (30-60%) of all cell lines tested, in a time- and concentration dependent manner. These results strengthen the idea of using 6-Br-Q0C10 to manage obesity. It is also implying that 6-BrQ0C10 may slow the growth rate of cancer cells and thus prolong life.

(This study was partially funded by NIH grants AA030221 and DA046258 to S.L.Chang.)

## Introduction

The energy required to sustain life, growth and the structural integrity of living organisms is derived mostly from a process named oxidative phosphorylation (1,2). This process takes place in the mitochondrial inner membrane of eukaryotic cells or in the cytoplasmic membrane of prokaryotic organisms, via a coupling reaction of two multi-subunit membrane protein complexes: the electron transfer chain complexes (Complexes I (3,4), II (5,6), III (7,8) and IV (9)) and ATP synthase complex (complex V) (10). The electron transfer chain complexes catalyze the oxidation of NADH, and succinate generated from TCA cycle in the catabolic oxidation of food staff, to generate proton gradient and membrane potential to drive the synthesis of ATP from ADP and inorganic phosphate, catalyzed by ATP synthase complex.

Among the four electron-transfer complexes, complexes I, III and IV generate proton gradient but complex II does not. Coenzyme Q_10_ (2,3-dimethoxy-5-(isoprenyl)_10_-6-methyl -1,4-benzoquinone) (11) (Figure 1A) mediates electron transfer between Complexes I or II and III. Coenzyme Q_10_ is the only redox component in the electron transfer system whose concentration can be manipulated by food intakes. The study of Coenzyme Q_10_ is extremely challenging, however, due to its low solubility. Fortunately, researchers (12,13) in the1970s have synthesized various coenzyme Q_10_ derivatives with shorter alkyl side chain to increase their solubility and yet still maintain their electron transfer activity. The electron transfer activity of these synthetic derivatives is dependent on the carbon chain length of the alkyl group of coenzyme Q, with the maximal activity reached with a length of ten carbon bonds. The most popular synthetic derivative is known as decyl-benzoquinone (DBQ, or Q_0_C_10_) (Figure 1B) (12). The spectral and redox properties of Q_0_C_10_ are identical to those of native coenzyme Q_10_ with an absorption maximum at 277nm and a midpoint redox potential of 110 mV. By replacing the 6-methyl group of the benzoquinone ring of Q_0_C_10_ with a bromine atom, our laboratory had synthesized 6-Br-decyl-benzoquinone (6-Br-Q_0_C_10_) (14). Figure 1C shows the chemical structure of 6-Br-Q_0_C_10_.

**Figure 1.**
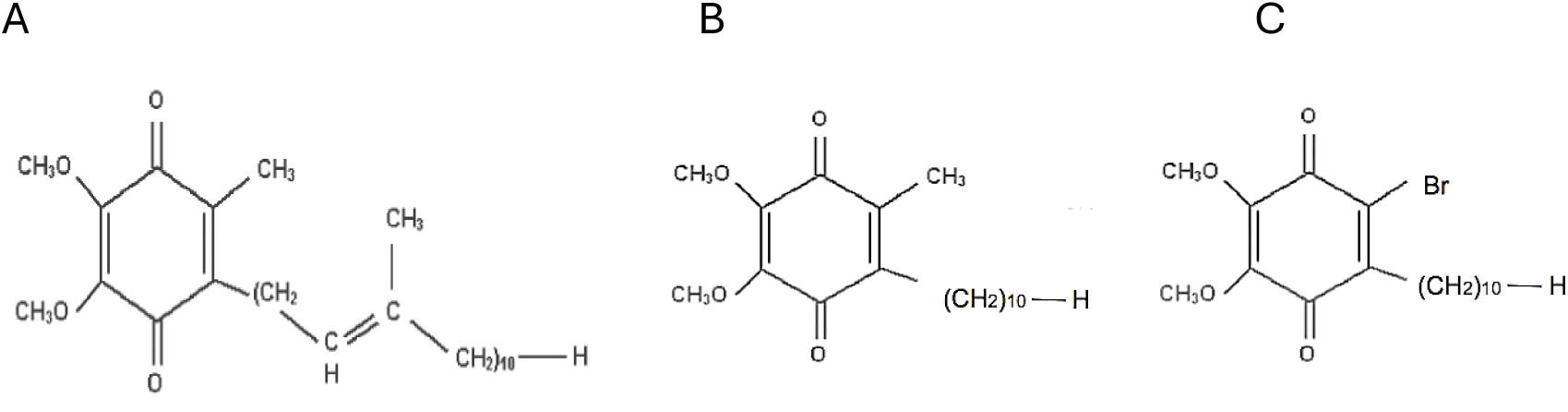
The chemical structure of coenzyme Q_10_ (2,3-dimethoxy-5-(isoprenyl)10-6-methyl-1,4-benzoquinone) (A), Q_0_C_10_ (2,3-dimethoxy-5-decyl-6-methyl-1,4-benzoquinone) (B), and 6-Br-Q_0_C_10_ (2,3-dimethoxy-6-decyl-6-Br-1,4 benzoquinone) (C).

Introducing an electron withdrawing group such as bromine (or halogen atom such as chlororine) should have an effect on the redox potential and the UV absorption property. Indeed, it has a higher redox potential than that of Coenzyme Q_10_, by about 30 mV and the UV absorption maximal peak redshift by 24 nm. A decrease of approximately 30% energy conservation efficiency has also been observed in isolated mitochondria that were treated with 6-Br-Q_0_C_10_. This decrease may be due to the loss of energy conservation site II at Complex III in the 6-Br-Q_0_C_10_ treated mitochondria.

Taking advantage of the availability of 6-Br-Q_0_C_10_, we have recently investigated the effect of this compound on cell growth using commercially available cell lines. As expected, the 6-Br-Q_0_C_10_ indeed reduces the cell growth by 30-60% depending on the type of cells used (15). These positive results indicate that 6-Br-Q_0_C_10_ (or 6-Cl-Q_0_C_10_) has great potential in reducing obesity, as it is caused by the oversupply of energy.

Although the results described in our recent report (15) suggest that 6-Br-Q_0_C_10_ has great potential to reduce obesity. it is obvious that more experiments on the effect of 6-Br-Q_0_C_10_ on animals and humans should be carried out before this approach can be further developed into useful remedies.

Recently, we have investigated the effect of 6-Br-Q_0_C_10_ on the growth of rats. When low concentration of 6-Br-Q_0_C_10_ (up to 0.38 mg/rat) was dissolved in different media, such as Carboxymethyl cellulose (CMC), ethanol, and mixture of oil and butter and fed to rats, they showed no toxicity and little effect on growth as measured body weight gain over a period. Rats that were given a higher concentration of 6-Br-Q0C10 (0.5 mg per rat) in a 0.3 mL mixture of oil and butter (66% oil, 33% butter) showed no increase in body weight over time. In contrast, rats that received only the carrier mixture experienced a 10.5% gain in body weight. Rats that received no treatment show an 8.6 % weight gain in the same period.

## Experimental

### Materials

6-Br-Q_0_C_10_: Crude 6-Br-Q_0_C_10_ was available in the laboratory (14). It was further purified to homogeneity using thin layer chromatography. The purity of 6-Br-Q_0_C_10_ was confirmed by its absorption peak at 303 nm and the concentration was determined spectrophotometrically, using a millimolar extinction coefficient of 20 cm^-1^(14).

Carboxymethyl cellulose (CMC) was obtained from Amazone. Oliver oil and unsalted butter were obtained from the grocery store.

#### Rats

Nine 3-week-old male F344 rats (born on 9/28/2023) were purchased from (Evigo RMS, Inc, Indianapolis, Indiana) on 10/17/2023.

### Methods

Three rats were housed in ventilated plastic cages (Animal Care Systems Inc., Centennial, CO). Rats were maintained in a temperature- and humidity-controlled environment with a 12-h light/dark cycle, provided with *ad libitum* access to a standard rat diet and water until 11/5/23. Rats were acclimated to the facility for 2 weeks before treatment.

The rats were divided into three groups (n=3): one group of rats was used for 6-Br-Q_0_C_10_ treatment, and the second group is used for treatment of carriers that used to dissolve 6-Br-Q_0_C_10_ and the third group was used as control without any treatment. All rats had free access to rat chow and water.

Various concentrations of 6-Br-Q_0_C_10_ were made in 1%. 1.5% CMC, 40% ethanol, and a mixture of oil and unsalted butter (66:33) (Table 1). 6-Br-Q_0_C_10_ solutions and carriers were given to rats via intragastric injection (i.g.) in a volume of 0.3 mL daily. The body weight of each rat was measured and recorded daily. The experimental protocol was approved by the Institutional Animal Care and Use Committee (IACUC) at Seton Hall University (IACUC Protocol No:SC2103).

**Table 1.** Doses of 6-Br-Q_0_C_10_ (mg/rat) and Carriers used.

| Dose # | Date | 6-Br-Q<sub>0</sub>C<sub>10</sub> (mg/rat) | Carrier |
| --- | --- | --- | --- |
| 1 | 11/5-11/11 | 0.015 | 1%CMC |
| 2 | 11/12-11/18 | 0.03 | 1%CMC |
| 3 | 11/19-11/25 | 0.06 | 1%CMC |
| 4 | 11/26-12/2 | 0.13 | 1.5%CMC |
| 5 | 12/3-12/27 | 0.23 | 1.5%CMC |
| 6 | 1/2-1/15 | 0.54 | 40%ethanol |
| 7 | 1/16-1/29 | 0.54 | Oil: butter (2:1) |

## Results

Figure 2 shows the body weight growth (weight gain) of rats from 11/5/2023 to1/16/2024 (Doses 1-7, 0.015 - 0.54 mg/rat) of all three groups, individually (Figure 2A) or as group average (Figure 2B). All of them showed a typical rat growth characteristic as from the vender’s report. There was no toxicity effect on the growth of rats, even if a high concentration of 6-Br-Q_0_C_10_ was used.

**Figure 2.**
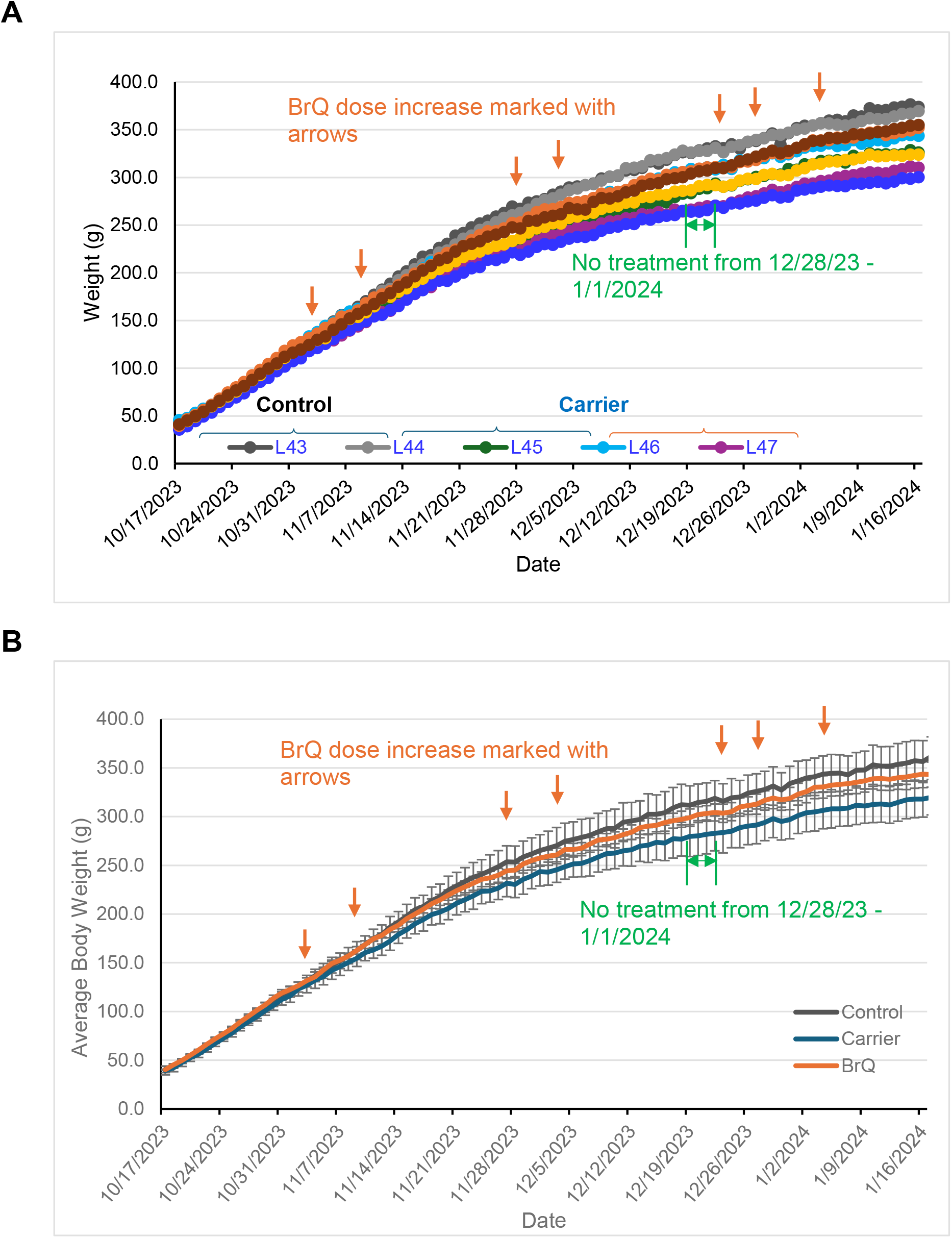
Assessment of Rat Body Weight Gain During 6-Br-Q_0_C_10_ Treatment. (A). Weight gain of all rats. (B). Group average weight gains of control, carrier and 6-Br-Q_0_C_10_-treated rats.

The amounts of 6-Br-Q_0_C_10_ used were from 30ug to 500ug per rat (see Table 1). The results indicated that 6-Br-Q_0_C_10_ treatment had no toxicity effect on rats even at high concentrations. The effect of 6-Br-Q_0_C_10_ on weight gain of rats was only slight when low concentrations of 6-Br-Q_0_C_10_ were used during the growing period of rats. A slight increase in food and water consumption was observed in the 6-Br-Q_0_C_10_ treated rats. The effect of 6-Br-Q_0_C_10_ on weight gain became more apparent when rats became more mature (age 61 to 71 days). When 6-Br-Q_0_C_10_ was used together with a high fat diet (the carrier), its effect on weight gain became very apparent. When higher concentration of 6-Br-Q_0_C_10_ (0.5mg/rat) was given to rats in 0.3 mL of oil/butter (66%/33%) mixture for a period, no body weight gains were observed, as compared to a 10.5% weight gain in the rats receiving the carrier mixture. Rats that received no treatment showed an 8.6 % weight gain in the same period. Figure 3 shows the effect of 6-Br-Q_0_C_10_ on weight gain when the high fat carrier was used.

**Figure 3.**
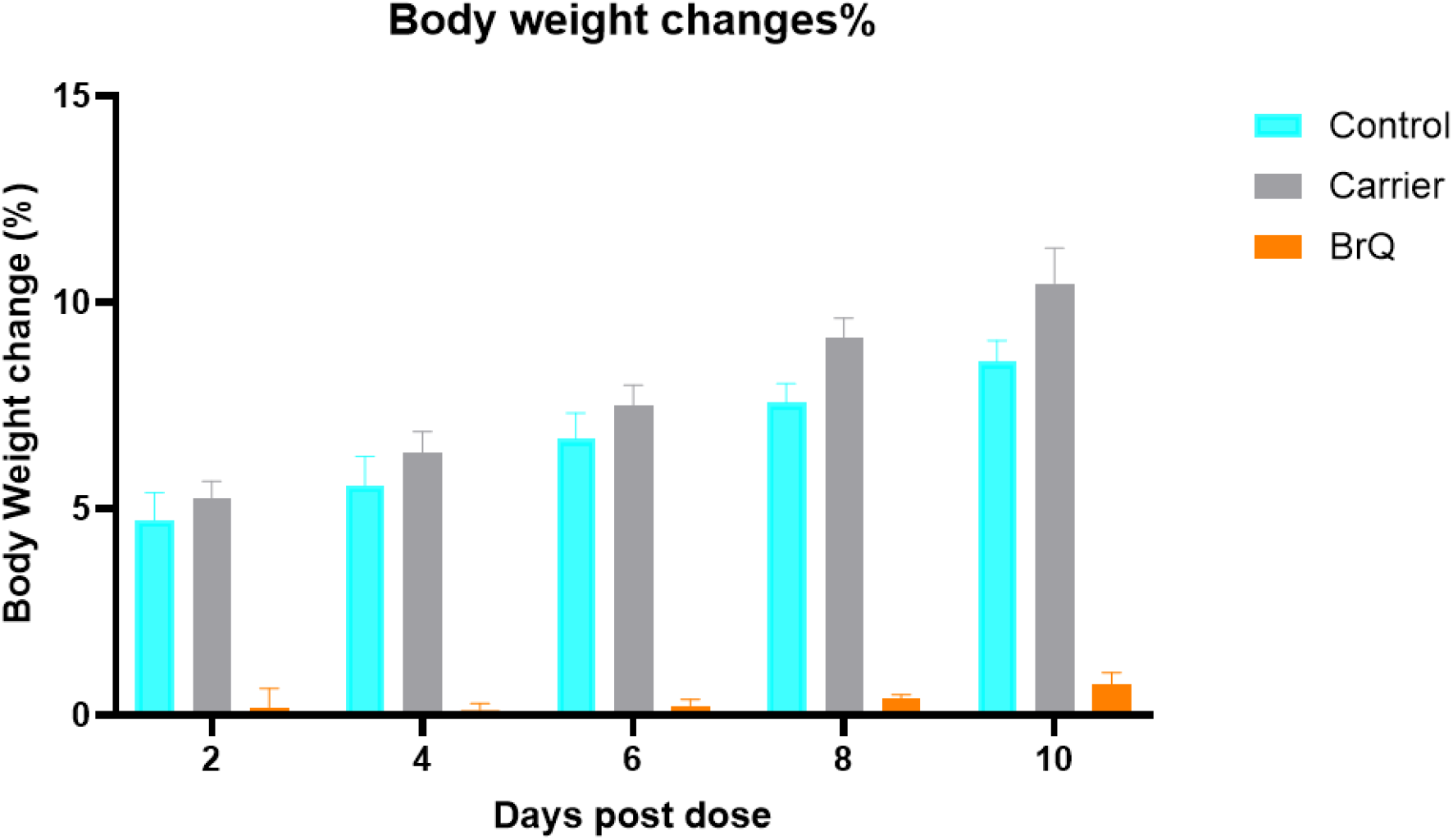
Effect on weight gains of rats fed with high-fat diet by 6-Br-Q_0_C_10_. Black (Control, normal diet), Blue (Carrier, high fat diet, 7 in Table 1) and Gold (high fat diet in the presence of 6-Br-Q_0_C_10_.

## Discussions

Animals obtain Coenzyme Q by two ways, *de novo* biosynthesis and absorption from food. It is known that the coenzyme Q_10_ content is higher in growing animals than in aged ones (16). Because the rats were fed with the same diet, the concentration of coenzyme Q_10_ in the growing rats must be resulted from biosynthesis. This inference is that 6-Br-Q_0_C_10_ is less effective in reducing weight gain in the growing rats than the mature ones, as the effectiveness has resulted from the competition between the 6-Br-Q_0_C_10_ and CoQ_10_, especially when a low concentration of 6-Br-Q_0_C_10_ is used. As expected, a more significant decrease in weight gain was observed in more mature rats with higher concentration of 6-Br-Q_0_C_10_.

When rats in their late period of growth were given 0.3 mL of high fat mixture daily in addition to a normal diet for ten days, a significant increase in weight gain was observed. This weight gain is diminished if 6-Br-Q_0_C_10_ is used in the mixture, indicating that the 6-Br-Q_0_C_10_ is replacing the normal Coenzyme Q and participating in the oxidative phosphorylation process in mitochondria of the treated rats. This result agrees with the observation that made in the 6-Br-Q_0_C_10_ treated isolated beef liver mitochondria (14). Using NADH linked substrate, 6-Br-Q_0_C_10_ treated mitochondria has a P/O ratio of 2 instead of 3. 6-Br-Q_0_C_10_ causes a bypassing of oxidative phosphorylation site 2 because of its low redox-potential.

The effect of 6-Br-Q_0_C_10_ on the reduction of weight gain might be enhanced, if it is used together with cholesterol lowing drugs such as Crestor or Lipitor, as they are also known to decrease the biosynthesis of coenzyme O_10_. Experiments on this aspect are currently in progress in our laboratory.

The obesity epidemic is one of the most serious problems that our country faces today (17,18). From a biological standpoint, obesity results from excess body fat accumulation, a condition which is primarily caused by the oversupply of energy generated from over consumption of foodstuff. Although a great number of specific diets aiming to reduce food intake are available, none of them are effective in curing obesity, because they are all based on the reduction of food intake, a restriction which obesity patients find difficult to accept. Although the most recent development in hormone-based drugs (GLP-1) offer a great opportunity in obesity management, they are by no mean perfect as they have many undesired side-effects and are of high cost. An alternative way of managing obesity is greatly needed. The use of 6-Br-Q_0_C_10_ or other low midpoint potential Q analogues to reduce energy recovery efficiency offers a great alternative in obesity management.

Also, it is conceivable that fast-growing cells (such as cancer cells) will be more affected by Br-Q_0_C_10_ than slower growing ones. Restricting the energy supply to cancer cells by the treatment of 6-Br-Q_0_C_10_ might slow down their growth and thus prolong the life of patients!

It is apparently metabolite of 6-Br-Q_0_C_10_ would be sodium bromide. It has a very weak toxic effect on biological system generally known as Bromism (19). The concentration used in this study, however, is too low to cause concern.

Other holo-Q_0_C_10_, such as 6-Cl-Q_0_C_10_, have been shown to be equally effective in reducing cell growth of many commercially available cell lines (15). It is highly likely that 6-Cl-Q_0_C_10_ will have the same effect of weight gain reduction and can be used in obesity management. Experimentation on this aspect is currently underway in our laboratory.

## Research ethical statement

The animal experimental protocol was approved by the Institutional Animal Care and Use Committee (IACUC) at Seton Hall University.

## Research funding

This study was partially funded by NIH grants AA030221 and DA046258 to SLChang.

